# Facial masculinity does not appear to be a condition-dependent male ornament in humans and does not reflect MHC heterozygosity

**DOI:** 10.1101/322255

**Authors:** Arslan A. Zaidi, Julie D. White, Brooke C. Mattern, Corey R. Liebowitz, David A. Puts, Peter Claes, Mark D. Shriver

**Affiliations:** Department of Biology; Pennsylvania State University, University Park, 16802, Pennsylvania, United States of America; Department of Anthropology, Pennsylvania State University, University Park, 16802, Pennsylvania, United States of America; Department of Electrical Engineering, ESAT/PSI, KU Leuven, Leuven, Belgium; Medical Imaging Research Center, MIRC, UZ Leuven, Leuven, Belgium

**Keywords:** facial masculinity, MHC, HLA, heterozygosity, height, sexual selection, human evolution, immunocompetence handicap hypothesis, ICHH, complex traits, condition-dependence

## Abstract

Facial masculinity is thought to be a condition-dependent male ornament, reflecting immunocompetence in humans. To test this hypothesis, we calculated an objective measure of facial masculinity/femininity using three-dimensional images in a large sample (N = 1,233) of people of European ancestry. We show that facial masculinity is positively correlated with adult height in both males and females. This suggests that variation in growth contributes, at least in part, to variation in facial masculinity, which is characteristic of condition-dependent traits. However, facial masculinity scales with growth similarly in males and females, suggesting that facial masculinity is not specifically a male ornament. Additionally, we measured immunocompetence via heterozygosity at the major histocompatibility complex (MHC), a well known genetic marker of immunity. We show that while height is positively correlated with MHC heterozygosity, facial masculinity is not. Thus, facial masculinity does not reflect immunocompetence measured by MHC heterozygosity in humans as thought previously. Overall, we find no support for the idea that facial masculinity is a condition-dependent male ornament that has evolved to indicate immunocompetence.

## Introduction

The immunocompetence handicap hypothesis (ICHH) (1–3) was proposed to explain the evolution of secondary sexual characteristics (*e.g.*, reindeer antlers and peacock trains), which grow to exaggerated proportions even though they might be detrimental to the fitness of the individual (4). According to the ICHH, androgens mediate the allocation of resources between the competing demands of fighting infection and the development of secondary sexual characteristics. Consequently, it is thought that males with more efficient immune systems can withstand greater immunosuppressive effects of androgens and can “afford” more extravagant displays (5–15). If this is true, then secondary sexual characteristics in males might be reliable (“honest”) indicators of physiological and immunological quality and strength to females and to other males (1, 2, 16). The ICHH has found some support in non-human animals (for review see Roberts et al., 2004 (17)). However, the mechanism through which sexual ornaments might have evolved to signal immunocompetence is not known.

A popular hypothesis that has been used to explain the evolution of sexual ornaments in non-human animals is condition-dependence (18–22). According to this hypothesis, sexual ornaments may have evolved to be more sensitive to the overall growth of the individual, which in turn is dependent on a variety of genetic and environmental factors, including immunocompetence, inbreeding, health status, and nutrient availability (23–25). Therefore, slight variation in immunocompetence among males could be amplified by large variation in the growth of sexual ornaments, making them ideal indicators of underlying health. Indeed, sexual ornaments are more variable in males and tend to be more sensitive to the overall growth of individual compared to other traits (23, 24, 26, 27). Traits such as tall height (28–30), facial masculinity (31, 32), and deep voices (33–37) are often thought of as sexual ornaments in humans as they are perceived to be attractive to females (38) and intimidating to other males (35, 39–41) However, the evidence that these traits reflect the condition or quality of males in humans is ambiguous and inconsistent across studies (17, 42–45). Some of the inconsistency could be due to methodological limitations of previous studies such as small sample size, lack of correction for population structure, and the use of perceived measures of masculinity and attractiveness, which are likely influenced by socio-cultural factors that are difficult to control in observational studies.

In this study, we investigated the ICHH in humans by testing for an association between immunocompetence, as measured by heterozygosity at the major histocompatibility (MHC) locus, and an objective measure of facial masculinity, a secondary sexual trait that is thought to have evolved to signal health and immune-function in humans (11, 46–50). Based on what we know about the condition-dependence of sexual ornaments (18–22), we tested three hypotheses:

#### Hypothesis 1

Facial masculinity is a condition-dependent male ornament in humans. If this is true, then we expect that facial masculinity would be a) correlated with height, b) more variable in males compared to females, and c) more strongly correlated with height in males compared to females.

#### Hypothesis 2

Immunocompetence is associated with adult height in humans. If immunocompetence plays a role in condition-dependent expression of secondary sexual characteristics, then it would be correlated with overall growth, measured by adult height, in humans.

#### Hypothesis 3

Facial masculinity reflects immunocompetence in men. Males who show greater immunocompetence should exhibit more masculine faces than males with lower immunocompetence, whereas facial masculinity should be less sensitive to variation in immunocompetence in females.

We used MHC heterozygosity to measure immunocompetence. The human MHC, also known as the Human Leukocyte Antigen (HLA) complex, is located on chromosome 6 and contains about 200 genes that are involved in immune-function (51). Higher genetic diversity at the MHC is correlated with immunocompetence, as it enables the immune system to recognize a more diverse array of foreign antigens (51–54). As a result, the MHC has experienced balancing selection in humans (51, 53–56) and therefore, heterozygosity at this locus serves as a useful proxy to measure immunocompetence.

In contrast with previous studies, we used an objective measure of facial masculinity, calculated at high resolution across the entire face, to test the ICHH in a large sample of persons of European ancestry. We also correct for genome-wide heterozygosity and population structure. Altogether our approach overcomes several key limitations of previous studies.

## Results

### Variation in facial masculinity

We calculated high resolution facial masculinity (FM) for the faces of 1,233 males and females of European ancestry from three-dimensional (3D) images, using a scalar-projection approach, similar to that described in (57) (Figure S6). 3D images were processed as described previously (58) allowing us to represent each face with a mesh of 7,150 points, or quasi-landmarks (QLs), each with *x*, *y*, and *z* coordinates (58, 59). For every QL in the face, the signed difference between the coordinates of the average female and male faces represents the direction of sexual dimorphism in 3D space (Fig. S6A). We define FM for each of the 7,150 QLs (FM_QL_) as the degree of change in a target face (X) along these vectors.

Fig. 1A shows a bimodal distribution of overall FM score (averaged across QLs - hereafter referred to as FM_overall_) where values of 0 and 1 represent FM_overall_ of the average female and male faces, respectively. The effect size of sex on FM_overall_ is comparable to that of height (Cohen’s D of 1.98 compared with 2.10 for height). The brow ridge, cheekbones, and nose ridge show the greatest degree of sexual dimorphism, in agreement with previous studies (Fig. 1B) (60).

**Fig. 1:**
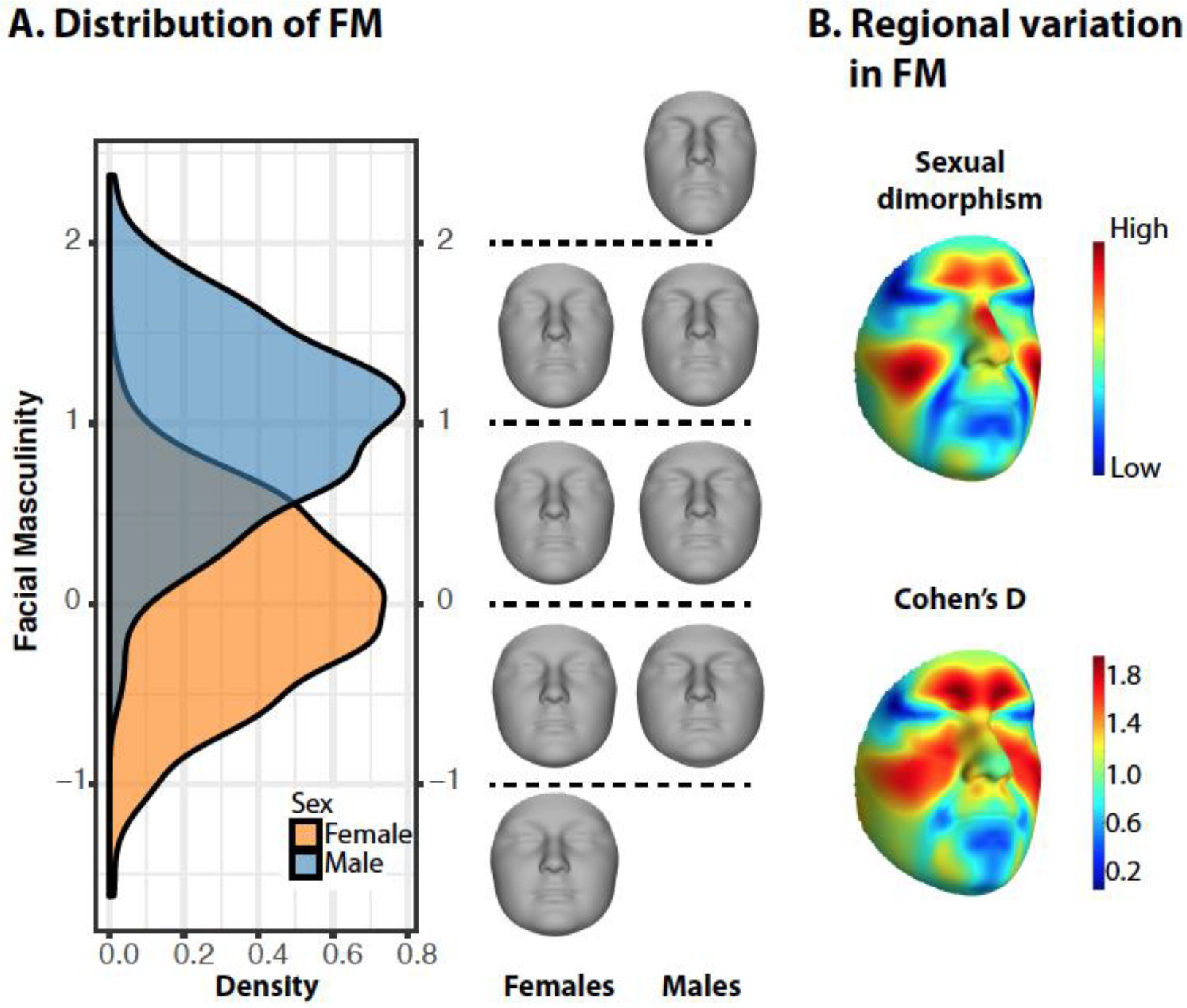
Sexual dimorphism in the face. A) Density plot showing a bimodal distribution of facial masculinity. B) Heatmaps showing the magnitude (top) and Cohen’s D (bottom) of sexual dimorphism between average male and female faces.

### Facial masculinity is positively correlated with height

We tested the relationship between overall facial masculinity (FM_overall_) and growth, using height as a predictor, with sex, age, weight, and genetic PCs 1-3 as covariates (Fig. S9). FM_overall_ is positively correlated with height, even within the sexes (Fig. 2A; Table 1; *t* = 8.81, *p* = 4.17 × 10^−18^), suggesting that taller people have more masculine faces than shorter people. Because variation in size of the faces was removed before calculating facial masculinity (Methods), this correlation represents allometric effects of growth on sexual dimorphism in face *shape*, not *size*. The effect of height on facial masculinity appears to be concentrated around the orbital region, nasal bridge, cheeks, and the chin, with masculinity in these regions increasing with height (Figure 2B). It is interesting to note that the distribution of the effect of height on masculinity across the face appears to be different from the effect of sex (Figure 2B).

**Fig. 2:**
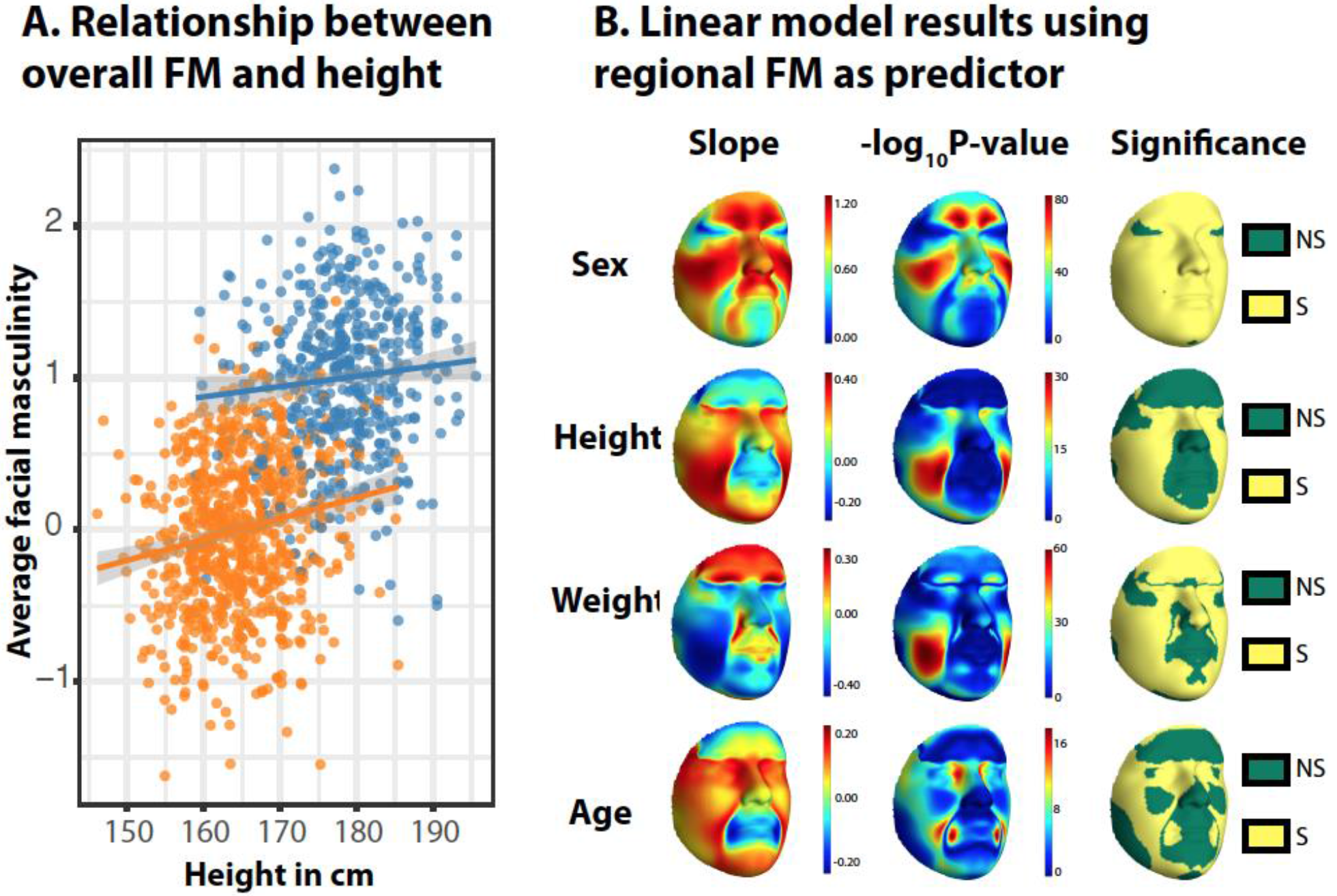
Facial masculinity is positively correlated with height, even within the sexes. A) Relationship between FM_overall_ and height. B) Results of linear model between FM_QL_ and height and other covariates. Regions in yellow are significant after Bonferroni correction for 7,150 QLs (*p* < 0.05/7150). Fig. S9 shows results for all covariates.

**Table 1:** Results of linear model between FM_overall_ and height with covariates. Slopes are standardized regression coefficients. Bonferroni cutoff for significance is 0.05 / 7 = 0.007.

|  | *Slope* | *SE* | *T statistic* | *P value* |
| --- | --- | --- | --- | --- |
| *Sex* | 1.240 | 0.058 | 21.46 | **$5.55 \times 10^{-87}$** |
| *Height* | 0.260 | 0.030 | 8.81 | **$4.17 \times 10^{-18}$** |
| *Age* | 0.109 | 0.020 | 5.60 | **$2.69 \times 10^{-08}$** |
| *Weight* | -0.221 | 0.023 | -9.64 | **$3.07 \times 10^{-21}$** |
| *gPC1* | -0.083 | 0.020 | -4.19 | **$2.94 \times 10^{-05}$** |
| *gPC2* | -0.063 | 0.019 | -3.25 | **$1.78 \times 10^{-03}$** |
| *gPC3* | -0.052 | 0.019 | -2.68 | $7.55 \times 10^{-03}$ |

The effect of height on FM_overall_ is not significantly different between males and females (*b*_*male*_= 0.227, *b*_*female*_ = 0.299, *Z-score*_*diff*_ = −1.18, *p* = 0.120), which agrees with the regional effects of height on FM_QL_ (Fig. S10). FM_overall_ is also not significantly more variable in one sex or the other (Levene’s test *p* = 0.675). Greater sensitivity to growth and higher variance in one sex relative to the other are classic signatures of sexual ornaments (19, 22, 24, 61), an expectation that facial masculinity does not meet. It is interesting to note that FM varies significantly along gPCs 1-3, suggesting that the patterns of sexual dimorphism and facial masculinity vary across populations, even within Europe (Table 1). This not only highlights the need to correct for population structure in future studies, but also calls for a detailed exploration of the variation in facial shape across populations.

### Height is positively correlated with immunocompetence

We fit a linear model between height and MHC heterozygosity, while correcting for genome-wide heterozygosity, sex, age, and gPCs 1-3. MHC heterozygosity shows a positive correlation with height (Table 2; *t* = 3.18, *p* = 0.0015), indicating that individuals who are more heterozygous at the MHC locus tend to be taller than people who are less heterozygous. This relationship is not driven by genome-wide heterozygosity, as the latter is not significantly associated with height (Table 2; *t* = −0.36, *p* = 0.72). Also note that height varies significantly along gPC1 and gPC2, which is consistent with the clinal variation in stature observed within Europe (62, 63)

**Table 2:** Results of linear model between height and MHC heterozygosity. Slopes are standardized regression coefficients. Bonferroni cutoff for significance is 0.05 / 7 = 0.007.

|  | *Slope* | *SE* | *T statistic* | *P value* |
| --- | --- | --- | --- | --- |
| *MHC het.* | 0.063 | 0.025 | 3.18 | **$1.54 \times 10^{-03}$** |
| *Genome-wide het.* | -0.007 | 0.020 | -0.36 | 0.72 |
| *Sex* | 1.452 | 0.040 | 36.20 | **$9.43 \times 10^{-196}$** |
| *Age* | 0.022 | 0.020 | 1.11 | 0.266 |
| *gPC1* | 0.122 | 0.020 | 6.11 | **$1.36 \times 10^{-09}$** |
| *gPC2* | 0.051 | 0.020 | 2.54 | **$1.14 \times 10^{-02}$** |
| *gPC3* | 0.014 | 0.020 | 0.70 | 0.49 |

### Facial masculinity is not correlated with immunocompetence

MHC heterozygosity is not significantly correlated with overall facial masculinity regardless of whether height is included in the model (*t* = −0.038, *p* = 0.970) or not (*t* = 0.586, *p* = 0.558). Thus, neither the allometric or non-allometric variation in facial masculinity is informative about MHC heterozygosity. Genome-wide heterozygosity is also not correlated with facial masculinity (height included as covariate: *t* = 0.02, *p* = 0.986; height not included as covariate: *t* = 0.65, *p* = 0. 516). MHC heterozygosity is also not significantly correlated with regional measures of facial masculinity (FM_QL_; Fig. S11). We also did not find any difference in the effect of MHC heterozygosity on facial masculinity between males and females (*b*_male_ = 0.035, *b*_*female*_ = −0.024, Z-score_*diff*_ = 0.32, *p* = 0.375; Fig. S12).

## Discussion

The immunocompetence handicap hypothesis (ICHH) has been widely used to explain the evolution of elaborate male ornaments. According to the ICHH, sexual ornaments are sensitive to variation in immune function, and signal immunocompetence and status to ‘discerning’ females and competing males. In humans, several sexually dimorphic traits such as deep voices, stature, strength, and facial masculinity have been regarded as male ornaments, based on the idea that these traits develop under the influence of androgens, which can be immunosuppressive, that women find these traits attractive (32, 64–66), and that other males find them intimidating (35, 39–41). We aimed to investigate whether facial masculinity is a male ornament in humans that indicates immunocompetence.

### Hypothesis 1: Facial masculinity is a condition-dependent male ornament in humans

Condition-dependent sexual ornaments tend to be highly variable and sensitive to variation in growth among males, which in turn can be dependent on a variety of genetic and environmental factors, such as immunocompetence, inbreeding, health status and nutrient availability (23–25). We find a significant positive correlation between height and facial masculinity in both males and females. Thus, overall growth appears to contribute, at least in part, to variation in facial masculinity among individuals. However, sexual dimorphism in the face is not simply due to an extended growth period in males compared to females, as evidenced from our observation that sex has a significant effect on facial masculinity, independent of height. Additionally, our results indicate that facial masculinity is not more variable in males compared to females and the pattern of correlation between height and facial masculinity is similar across the sexes. Taken together, these results fail to support the hypothesis that facial masculinity is a condition-dependent male ornament, which should be more variable and more sensitive to growth in males (24, 25).

### Hypothesis 2: Immunocompetence influences growth in humans

There is a positive correlation between height and MHC heterozygosity, but not at loci across the rest of the genome. This result is consistent with the positive relationship between height and other measures of immunocompetence such as antibody titers (67, 68).

### Hypothesis 3: Facial masculinity is an expression of immunocompetence

Facial masculinity is not correlated with MHC heterozygosity, suggesting that facial masculinity does not reliably indicate this measure of immunocompetence. We also find no support for the contention that FM indicates heterozygosity across the genome generally, something that has also been proposed previously (20, 69).

Altogether, our results call into question some of the evolutionary explanations behind female and male perceptions of facial masculinity, and whether facial masculinity should be regarded as a condition-dependent sexual ornament in humans. Yet, our data show that sex differences in face shape are not merely developmental byproducts of greater male size. Little is known about the genetic and environmental factors contributing to differences in facial shape between males and females, or the variation in facial masculinity within the sexes. Differences in facial shape exist between male and female children as young as three years old (70, 71), and are likely defined, in part, during gestation by the intrauterine environment (72–74). Sexual dimorphism in the face increases dramatically at the onset of puberty, implicating sex hormones and other endocrine processes underlying general growth occurring around this period (70, 75–79). We do not know whether this degree of sexual dimorphism in facial shape is new to humans since their divergence from other hominins and apes, and if it perhaps evolved as a mechanism to intimidate rival males (80–82). Alternatively, facial sexual dimorphism may represent a vestigial trait that has decreased over time as a result of self-domestication (83–85). We also do not know how facial sexual dimorphism varies across human populations. We suspect that facial sexual dimorphism varies considerably across human populations, both in degree and pattern. We show that facial masculinity varies across populations, even within Europe. Is genetic drift sufficient to explain these patterns? How much have differences in perceptions of beauty and social status across populations shaped the evolution of sexual dimorphism, and facial shape in general? These gaps in our knowledge highlight the need for a more mechanistic understanding of both genetic and environmental factors underlying the development of sexual dimorphism, as well as an appreciation of the variation in facial shape within and across diverse human populations before we can cultivate a clearer picture of the role of sexual selection in human evolution.

## Materials and Methods

### Participant recruitment

Study participants were recruited in the United States through the Anthropology, DNA, and the Appearance and Perceptions of Traits (ADAPT) Study based at The Pennsylvania State University under Institutional Review Board (IRB) approved protocols (#44929, #45727). 3D images were taken using the 3dMD Face system (3dMD, Atlanta, GA). Height and weight were measured using an Accustat stadiometer (Genentech, South San Francisco, CA) and clinical scale (Tanita, Arlington Heights, IL). Genotyping was conducted by 23andMe (23andMe, Mountain View, CA) on the v4 genome-wide SNP array. After filtering out SNPs with more than 10% missing genotypes, this array comprised of approximately 600K SNPs.

### Data curation

From the 2,721 participants with faces and genotype data, we removed individuals with missing covariate data, misclassified sex information, and individuals with more than 10% missing genotypes. We further restricted the analysis to unrelated individuals between 18 years of age and 30 years of age (to reduce the effects of aging). Relatives were identified as pairs of individuals with an identity-by-state (IBS) value of at least 0.8, after which one of each pair was removed, resulting in a set of 1,921 unrelated individuals (Fig. S1).

### Ancestry and population structure

We selected people of European ancestry as they were the largest sample in our dataset. To do so, we merged the genotype data from our sample (N = 1,921) with genotypes from the 1,000 Genomes Project dataset (N = 2,503). Prior to merging the genotype data, we removed SNPs that did not intersect between the two datasets, palindromic (A/T, G/C) SNPs, and SNPs that did not meet standard quality-control criteria (Fig. S1). SNPs were further pruned for linkage disequilibrium with a window size of 50 SNPs, a step size of 5 SNPs, and a variance inflation factor threshold of 2 using PLINK 1.9 (86, 87), resulting in 201,042 SNPs. Genetic ancestry was inferred using an unsupervised clustering scheme in ADMIXTURE with K ranging from 2 to 16 (Fig. S2 & S4) (88). We selected results from K=6 as this value had a low cross-validation error (Fig. S3) and showed separation based on continental ancestry (Fig. S2). 1,249 individuals of primarily European ancestry were identified based on ADMIXTURE output in comparison with European samples from the 1,000 Genomes Project (Fig. S4). We carried out principal components analysis on the genotypes of this subset and removed 16 outliers using the smartpca program in Eigensoft (Fig. S5) (89, 90), leading to a sample size of 1,233 individuals. The first three genetic PCs (gPCs) were used as covariates to correct for population structure.

### Processing 3D photographs

High-resolution 3D images were ‘cleaned’ to remove hair, ears, and disassociated polygons. Five positioning landmarks were placed (two on the inner corner of the eyes, two on the outer corners of the mouth, and one on the tip of the nose) to establish facial orientation. An anthropometric mask comprised of 10,000 quasi-landmarks (QLs), which is later trimmed to 7.150 QLs, was non-rigidly mapped (91) onto all 3D surfaces such that each QL is spatially homologous across individuals (92). Thus, every face can be represented by a configuration of 7.150 QLs, each with three coordinates (x, y, and z). For each face, a mirror image was created by changing the sign of the x coordinates following (93), which was mapped with QLs in the same way as the original, non-reflected face. A generalized Procrustes superimposition (94) of both the original and reflected images together was performed to eliminate differences in position, orientation, and scale. At this point, the variation in face shape can be decomposed into the symmetric (average of mirror QL configuration) and asymmetric (difference of mirror QL configurations) components of facial shape (60).

### Calculating facial masculinity

We define facial masculinity/femininity as the magnitude of change in aspects of the face that are different, on average, between males and females in the population. Thus, facial masculinity can be represented as the degree of change in the direction from an average female face to an average male face. Conversely, facial femininity is the magnitude of change in the opposite direction. We calculated facial masculinity (FM) per quasi-landmark (QL) for every face, using a scalar-projection approach (Figure S6) (57). First, we generate female and male consensus faces from the sample by averaging the QL configurations across all females and all males, respectively. For every QL in the face, the signed difference between the coordinates of the male and female consensus faces is a three-dimensional vector 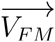 that represents the direction of sexual dimorphism in 3D space (Figure S6A). The goal is to calculate the degree of change in each QL of a target face X along these vectors (*i.e.,* one for each of the 7,150 QLs), which is the FM per QL (FM_QL_). This can be done by computing the scalar projection of 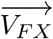 the difference between X and the female consensus face, onto 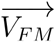 (Figure S6C).

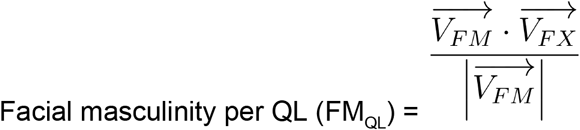

### Genomic and MHC heterozygosity

We defined individual heterozygosity as the proportion of heterozygous SNPs in a region. Genome-wide heterozygosity was calculated from a total of 192,417 LD-pruned, autosomal SNPs. To measure MHC heterozygosity, we obtained a list of 195 SNPs tagging haplotype variation for the classical HLA genes in Europeans (95). We used 114 of these SNPs, the subset for which our samples were genotyped (Fig. S7), to calculate MHC heterozygosity. These SNPs capture most of the HLA alleles (95) and the heterozygosity calculated using the subset of 114 SNPs is highly correlated with heterozygosity calculated using all 195 markers in the sample of Europeans available in the 1000 Genomes Project dataset (Fig. S8) (96).

### Data availability

The informed consent with which the data were collected does not allow for dissemination of identifiable data to persons not listed as researchers on the IRB protocol. Thus, the raw genotype data and 3D images cannot be made publically available. In the interest of reproducibility, we have provided de-identified overall facial masculinity measures as well as age, sex, weight, height, ancestry, genetic PCs, and MHC and genome-wide heterozygosity, from which all results presented in this manuscript can be reproduced. In addition, we also provide high-density facial masculinity maps, i.e, facial masculinity calculated for every quasi-landmark (FM_QL_) for all 1,233 individuals used in the analyses. This resource will allow other researchers to study variation in facial masculinity with high resolution in a large sample. We have made all scripts available on the following GitHub repository: https://github.com/Arslan-Zaidi.

## Acknowledgements

We thank the participants for providing the data necessary to carry out this study. We are grateful to the members of Shriver Lab and Puts Lab for helping with data collection. Finally, we would like to thank Tina Lasisi and Tomás González-Zarzar for helpful discussions on the manuscript.

